# Endothelial-derived exosomes demonstrate a link between endothelial innate inflammation and brain dysfunction and injury in aging

**DOI:** 10.1101/670083

**Authors:** F.M. Elahi, D. Harvey, M. Altendahl, K.B. Casaletto, N. Fernandes, A.M. Staffaroni, P. Maillard, J.D. Hinman, B.L. Miller, C. DeCarli, J.H. Kramer, E.J. Goetzl

**Author notes:** Corresponding Authors: Fanny M. Elahi, MD PhD, 675 Nelson Rising Lane, Suite 190, San Francisco, CA, 94158, Edward J. Goetzl, MD, Geriatric Research Center, 1719 Broderick St., San Francisco, CA 94115.

## Abstract

We test the hypothesis that endothelial cells take on an inflammatory phenotype in functionally intact human subjects with radiographic evidence of white matter injury. Markers within all three complement effector pathways and regulatory proteins were quantified from endothelial-derived exosomes (EDE) of subjects (age 70-82) with (n=11) and without (n=16) evidence of white matter hyperintensity on MRI. Group differences and associations with systemic markers of immune activation (IL6, ICAM1), cognition and neuroimaging were calculated via regression modelling.

EDE complement factors within the alternative and classical pathways were found to be higher and regulatory proteins lower in subjects with WMH. EDE levels of several factors demonstrated significant associations with cognitive slowing and systolic blood pressure. The inhibitor of the membrane attack complex, CD46, showed a significant positive association with cerebral grey matter volume. Systemic inflammatory markers, IL6 and ICAM1, were positively associated with EDE levels of several factors.

These findings provide the first *in vivo* evidence of the association of endothelial cell inflammation with white matter injury, cognition, and brain degeneration in functionally normal older individuals, and form the basis for future biomarker development in early or preclinical stages of vascular cognitive impairment and dementia.

## 1. INTRODUCTION

Cerebrovascular disease and the associated blood-brain barrier (BBB) dysfunction are intimately associated with immune activation and among the most common age-associated, inflammation-mediated, degenerative brain changes. Vascular pathways are emerging as an important contributor to neurodegenerative disorders^1–4^. Importantly, immuno-vascular dysregulation can cause pathological systemic-brain cross-talk^5^ in early disease states, prior to frank brain degeneration and clinical manifestations such as mild cognitive impairment^6,7^. More recently, molecular pathways are emerging to suggest a feed-forward degenerative-inflammatory phenomenon between endothelial cells, innate immune activation, and degenerative myelin debris^8^. Therefore, identification of molecular biomarkers of immuno-vascular disease in preclinical states has important therapeutic implications for extension of health span, treatment of vascular cognitive impairment and associated neurodegenerative processes, such as Alzheimer’s disease^9^. However, detection of early or preclinical cellular dysfunctions has been challenging.

In light of the inaccessibility of brain cells, neuroimaging techniques have been developed to capture indirect consequences of cerebrovascular disease (CVD), such as white matter hyperintensities (WMH) on T2/Flair (fluid-attenuated inversion recovery) imaging. However, imaging alone does not suffice, as the underlying molecular etiology of radiographic white matter changes^10^ can be diverse in aging and across neurodegenerative disorders. Exosomal cargo can provide molecular information on disease processes affecting inaccessible organs, including the brain ^11–16^. Exosomes are plasma membrane- and endosomal-derived vesicles, that get released by most cell types, and carry cargo molecules from their cell of origin including proteins, mRNAs, microRNA and lipids ^17,18^. Cerebral cell-derived exosomes range in diameter from 30 nm to 220 nm and can readily cross the blood-brain barrier ^19^. Analyses of exosome-derived molecular cargo are providing an unprecedented opportunity to non-invasively investigate cell-specific pathological change *in vivo*, with great impact on diagnostics of diseases affecting inaccessible organs ^20^.

In the aging brain, WMH on MRI can frequently be seen preclinically and presumed secondary to immuno-vascular effects, involving injury of endothelial cells (ECs) at the blood-brain barrier and increased CNS immune activation and reactive gliosis^21^. Employing a precision medicine approach in functionally-intact subjects, we investigated concentrations of complement factors of innate immunity derived from endothelial-derived exosomes (EDEs) isolated from plasma to provide the first *in vivo* test of the hypothesis that endothelial cells take on an inflammatory phenotype in association with radiographic evidence of WMH.

Complement factors, essential for innate immunity, form a collection of more than thirty soluble proteins that work with leukocytes to protect the host from pathogens. Complement factors are divided into three distinct but highly connected pathways: the classical, the alternative, and the lectin pathways, as well as complement regulatory proteins (**Figure 1**). When dysregulated, complement activation can cause robust destruction of host cells. Complement factors also increase cytokine and chemokine production, amplifying inflammation and leading to activation and recruitment of immune cells^22,23^. Interactions between inflammatory cytokines and EC complement proteins may enable inflammatory homeostasis or exacerbate a dysfunctional state. Mammalian ECs in culture have been shown to produce many complement proteins including C1, C4, C3, factor B, and factors I and H^24^.

**Figure 1.**
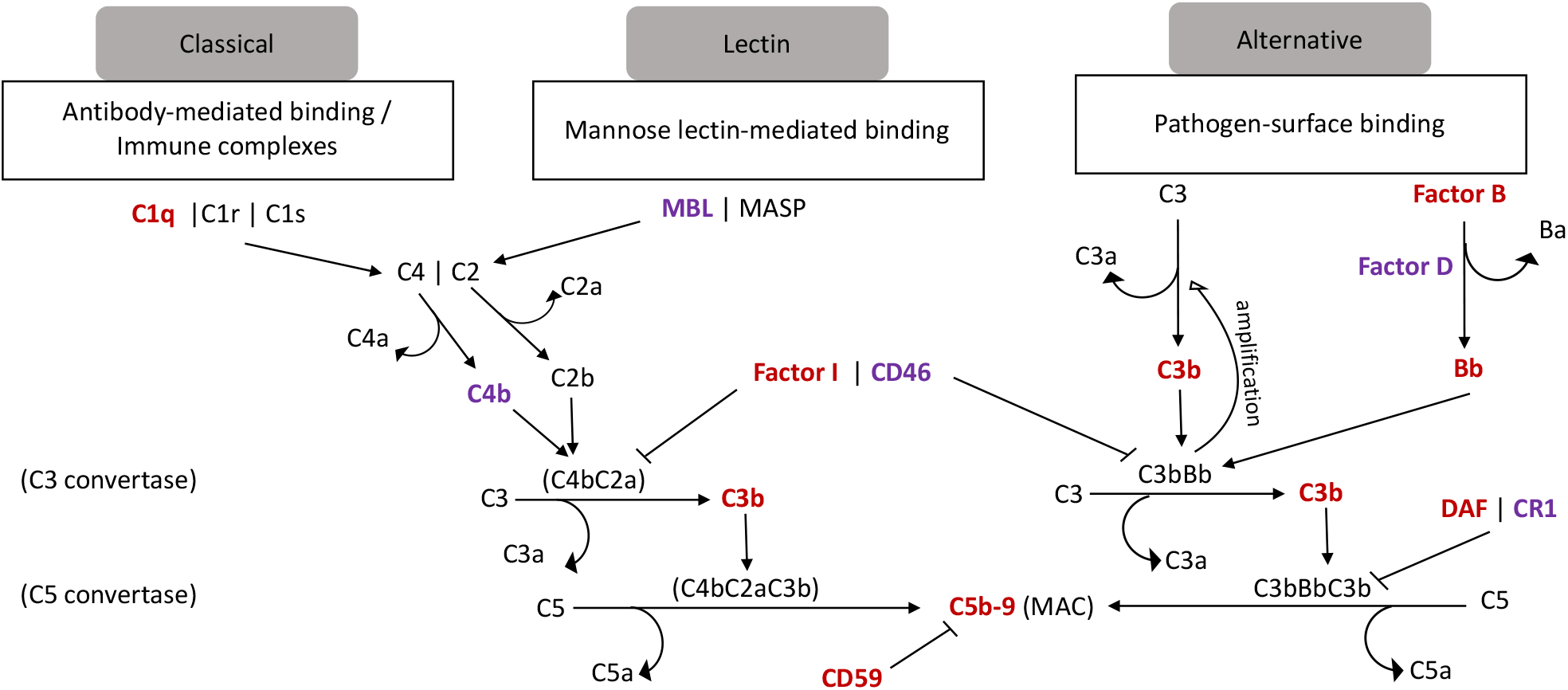
Complement Pathways. In red bold complement factors with significantly different levels between groups, and in purple bold complement factors not found to have significantly different levels between groups. The extreme inflammatory consequences of a deficiency of a single complement regulatory protein emphasizes the largely non-redundant organization of complement control mechanisms. Once complement proteins are secreted by ECs or recruited from other cells to the surface of ECs, the classical complement effector system may be activated by several mechanisms. The binding of C1q to EC C1q receptors is a minor complement activation pathway, but the greater respective roles of C1q bound to EC surface C-Reactive Protein, pentraxin 3, phosphatidylserine or to antibodies on proteins such as class I HLA remain to be definitively elucidated. C5b-9 MAC is capable of injuring ECs directly by inducing membrane damage, whereas C3b injures ECs indirectly by attracting monocytes, enhancing their adherence to ECs and stimulating their diverse cytotoxic mechanisms.

At a time when the goal of therapies is the gain specificity, a major hurdle to progress has been the low specificity of biomarkers for endothelial contributions to BBB dysfunction and neurodegeneration. To this end, EDE biomarkers provide the opportunity to specifically interrogate endothelial molecular and quantify their associations with vascular risk factors as well as downstream pathological changes. We present a proof of concept study, where we investigate levels of complement factors proteins and their regulatory proteins in endothelial cells of functionally normal subjects with WMH by using EDE cargo analyses as a means of performing a “liquid biopsy” of inaccessible endothelial cells *in vivo*. We then investigate the association of EDE complement factors with an important vascular risk factor, blood pressure, as well as downstream changes to brain structure and function.

## METHODS

### 2.1. Study Participants

The study cohort and sampling have been previously described in detail^25^. In brief, we minimized bias by performing a consecutive sampling of functionally intact, older study participants from ongoing longitudinal studies of brain aging at the Memory and Aging Center at UCSF (NIH Aging and Cognition study; Larry J. Hillblom foundation study; NIH Chronic Inflammation study) to prospectively participate in this study. The characteristics of subjects have been provided in detail in our prior publication using this cohort^25^. The main inclusion criteria was presence or absence of white matter injury on brain MRI. A total of 26 subjects were enrolled, with 11 included in the cerebral small vessel disease (cSVD) group, classified based on global cerebral volume of WMH on T2/FLAIR (modified Fazekas score of >=2), and 15 controls who had no significant WMH on brain MRI (Fazekas=0). The main exclusion criteria were confounding neurological or psychiatric disease, strokes, brain tumors, and inability to safely obtain an MRI. All brain MRIs were rated for burden of WMH by a board-certified neurologist (FME). At the time of enrollment in the current study platelet-poor-plasma was prepared according to published methods^25,26^. Plasma samples were aliquoted and stored at −80°C. All study participants provided informed consent and the study protocols were approved by the UCSF Human Research Protection Program and Institutional Review Board. Research was performed in accordance with the Code of Ethics of the World Medical Association.

### 2.2. Cognition (Processing Speed)

As previously detailed^25^ all study participants completed a modified version of the Trail-Making Test, that was used as a measure of speeded set-shifting^27^, which is thought to be affected by cSVD. Briefly, this timed task requires participants to sequentially alternate between numbers and days of the week as quickly as possible. The variable used in analyses reported in this work is the time taken to perform the task (log transformed to normalize the distribution).

### 2.3. Neuroimaging Evaluation

#### MRI Acquisition

All participants were scanned on a Siemens Prisma 3T scanner at the UCSF Neuroscience Imaging Center. A T1-weighted Magnetization-prepared rapid gradient echo (MP-RAGE) structural scan was acquired in a sagittal orientation, a field-of-view of 256 x 240 x 160 mm with an isotropic voxel resolution of 1 mm3, TR = 2300 ms, TE = 2.9 ms, TI = 900 ms, flip angle = 9°. The T2 fluid attenuated inversion recovery (FLAIR) acquired in the axial orientation, field-of-view = 176 x 256 x 256 mm, resolution 1.00 x 0.98 x 0.98 mm3, TR = 5000 ms, TE = 397 ms, TI = 1800 ms.

#### MRI Processing and Analyses

De-identified digital information was transferred from UCSF using secure and HIPAA complaint DICOM server technology Images were processed by the Imaging of Dementia and Aging (IDeA) lab at UC Davis and full imaging protocol details are reported in prior publications^28–31^. In brief, WMH quantification was performed on a combination of FLAIR and 3D T1 images using a modified Bayesian probability structure based on a previously published method of histogram fitting. Prior probability maps for WMH were created from more than 700 individuals with semi-automatic detection of WMH followed by manual editing. Likelihood estimates of the native image were calculated through histogram segmentation and thresholding. All segmentation was initially performed in standard space resulting in probability likelihood values of WMH at each voxel in the white matter. These probabilities were then thresholded at 3.5 SD above the mean to create a binary WMH mask. Further segmentation was based on a modified Bayesian approach that combines image likelihood estimates, spatial priors, and tissue class constraints. The segmented WMH masks were then back-transformed on to native space for tissue volume calculation. Volumes were log-transformed to normalize population variance.

### 2.4. Enrichment of Plasma EDEs and Extraction of Cargo Proteins

Platelet-poor plasma was prepared from 6 ml of venous blood and stored in 0.5 ml aliquots at −80°C as previously described ^26^ and EDE were enriched as per previously published protocol^26^. Briefly, after depletion of platelets, EDE exosomes were enriched by sequential immunoprecipitation with two biotinylated monoclonal antibodies to CD31 (MEM-05, Thermo Fisher Scientific) and then CD146 (Novus Biologicals, Littleton, CO, USA) prior to lysis of exosomes for quantification of cargo proteins via ELISA. Sequential rounds of immunoprecipitation aimed to enhance selectivity. According to consensus criteria for characterization of extracellular vesicles^32^, we used Nanosight LM10 instrument (Malvern Instruments, Malvern, UK) to confirm that the particle sizes of total exosomal extracts were within range for exosomes (30-220 nm) in our prior studies^25,26,33^. We used the exact same technique for the work being reported in this manuscript and therefore we are making the assumption that the immunoprecipitated particles will also be within size range.

### 2.5. Enzyme-Linked Immunosorbent Assay Quantification of Proteins

Commercially available colorimetric ELISA kits were used for the quantification of exosomal proteins, including tetraspanin exosomal marker CD81 (Cusabio; American Research Products, Waltham, MA, USA), complement fragment C4b (Cusabio Technology, College Park, MD), complement receptor 1, and decay accelerating factor (CR1, DAF; ARP American Research Products, Waltham, MA), factor I (Cloud-Clone Corp, Katy, TX), complement fragment C3d and CD46 (LifeSpan Biosciences, Seattle, WA), complement fragment C3b, Factor B, C1q portion of the C1 complement complex (Abcam, Cambridge, MA), Bb fragment of complement factor B (Quidel-Microvue, San Diego, CA), terminal complement complex C5b-C9 (Elabscience, Bethesda, MD), CD59 and mannose-binding lectin (MBL; Ray Biotech, Norcross, GA), and complement factor D (ThermoFisher-Invitrogen, LaFayette, CO). The mean value for all determinations of CD81 in each assay group was set at 1.00, and relative values of CD81 for each sample were used to normalize their recovery. As per our prior work^25,26^ and that of others, CD81 was used as a surrogate marker of exosome concentration against which each protein was normalized.

### 2.6. Quantification of Plasma Analytes

As previously described^25^, plasma cytokine concentrations were measured by high-performance electrochemiluminiscence (HPE) using the validated, commercially available multiplex V-PLEX Human Proinflammatory (IL-6) and Human Vascular Injury (ICAM1) assays. The analytes, IL-6 and ICAM1 were selected to reflect global levels of systematic inflammation and immune activation. The multiplex arrays were analyzed with a MESO QuickPlex SQ 120 imager (MSD, Rockville, MD) and Discovery Workbench v4.0 software. Each sample concentration was measured in duplicate in accordance with the manufacturer’s protocol.

### 2.7. Statistical Analyses

JMP Pro and PRISMA softwares were used for statistical analyses. Significantly skewed variables were log transformed to normalize distributions. Analyses included chi-squared tests (for categorical variables) and Student’s t-tests (for continuous measures). Linear models were additionally conducted to determine group EDE differences, adjusting for age, and discriminant analyses were employed for ROC analyses to determine group classification accuracy of EDE biomarkers. Finally, linear models examined associations between EDE levels with plasma inflammatory markers, blood pressure, cognitive speed, and structural brain MRI outcomes, adjusting for age and TIV, as appropriate. We also performed correction for multiple comparisons using the Benjamini-Hochberg False Discovery Rate (FDR)^34^. For illustration of results, all p-values were rounded to one non-zero digit beyond the decimal.

## 3. RESULTS

### 3.1. Demographics (Table 1, Figure 2)

Numerical values are summarized in **Table 1**. The mean age for subjects with WMH was significantly higher than controls (p=0003). We therefore controlled for age in all our analyses. There were no significant differences with respect to sex and educational attainment between groups. The global measure of cognitive function, mini-mental status exam (MMSE), did not significantly differ between groups (p=.60). With respect to vascular risk factors, subjects with WMH had on average higher systolic blood pressures (p=.0005). No significant difference was noted for diastolic blood pressure (p=.70). Diagnosis of hypercholesterolemia (p=.90), or hyperglycemia (p=.30) did not differ between groups, nor were any significant differences found in laboratory values for fasting lipid panel (triglycerides p=.40; HDL p=.40; and LDL p=.10), blood insulin concentration (p=.90), hemoglobin-A1C (p=.40), and the homeostatic model assessment of insulin resistance (HOMA-IR) (p=.60)—a surrogate for assessing β-cell function.

**Figure 2.**
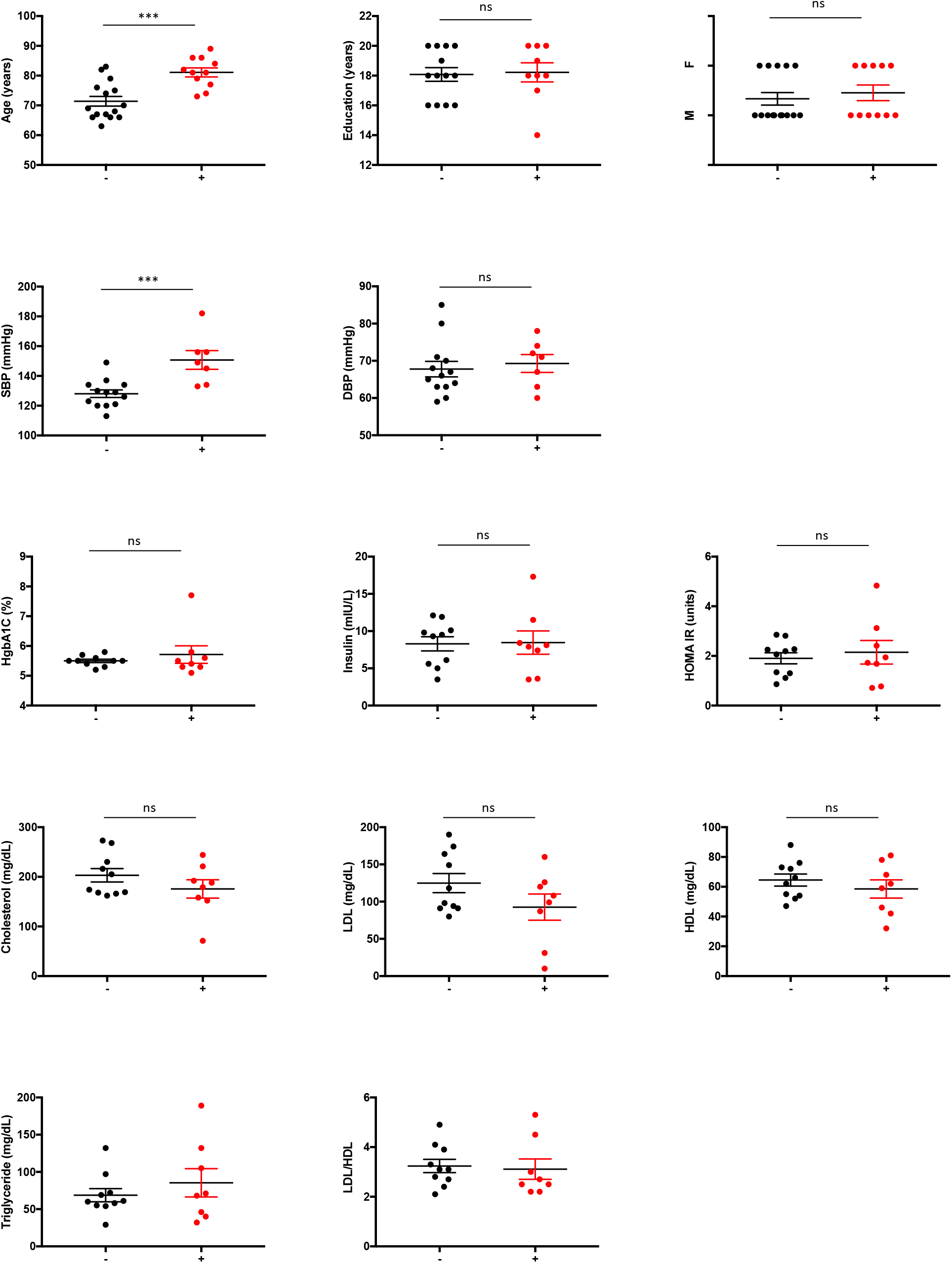
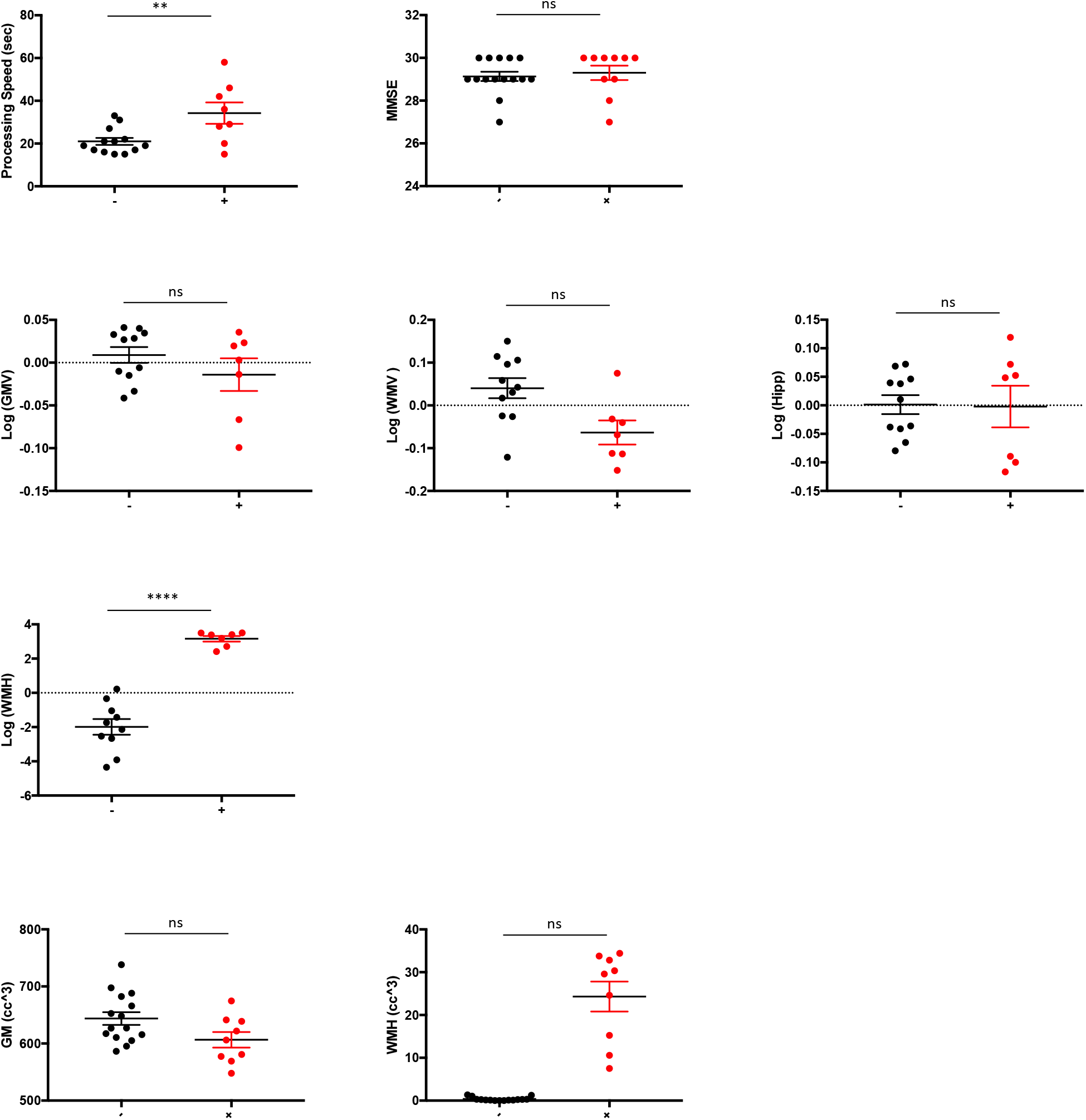
Demographic, cognitive and vascular risk group comparisons between subjects with WMH and without. (A) Demographics include age, gender, and education; (B) vascular risk factors include systolic and diastolic blood pressure, as well as (C) clinical labs: hemoglobin A1C, insulin, homeostatic model assessment of insulin resistance, total cholesterol, low density lipoprotein, ratio of low to high density lipoprotein, and triglycerides; (D) neuroimaing: white matter hyperintensity, global white matter volumes, global grey matter volumes, and mean of bilateral hippocampal volumes. For volumetric neuroimaging measures, total intracranial volumes were regressed out and residual used in models. Dependent variables were log transformed as needed to approximate normal distributions. All significance levels are from ANCOVA for continuous variables controlling for age, and Chi-square for gender. Grey matter and white matter hyperintensity volumes are also shown as raw values. Acronyms: SBP=systolic blood pressure; DBP=diastolic blood pressure; HgbA1c=hemoglobin A1C; HOMA-IR= homeostatic model assessment of insulin resistance; LDL=low density lipoprotein; HDL=high density lipoprotein; MMSE=mini-mental status examination; WMH=white matter hyperintensity; WMV=white matter volume; GMV=grey matter volume.

**TABLE 1.** Participant demographic, cognitive, and clinical data. Two-tailed Student’s t-tests or the non-parametric Wilcoxon rank sum were used for statistical comparison of continuous measures and Chi-squared tests were performed to compare group characteristics. Results are included in this table. Age, systolic blood pressure, and processing speed were the only significant different variables between groups.

|  | WMH- | WMH+ | p value |
| --- | --- | --- | --- |
| Total Number | 15 | 11 | - |
| Female, % total (N) | 40 (6) | 45 (5) | 0.50 |
| Age, Mean years (SEM) | **72 (2)** | **80 (2)** | **0.0003** |
| Education, Mean years (SEM) | 18 (.5) | 18 (.6) | 0.90 |
| CDR | 0 | 0 | - |
| MMSE, Mean (SEM) | 29 (.2) | 29 (.3) | 0.60 |
| Processing Speed, seconds (SEM) | **21(6)** | **34 (14)** | **0.007** |
| Systolic Blood Pressure, mmHg (SEM) | **130 (3)** | **150 (6)** | **0.0005** |
| Diastolic Blood Pressure, mmHg (SEM) | 68 (2) | 69 (2) | 0.70 |
| Low Density Lipoprotein, mg/dL (SEM) | 125 (12) | 93 (18) | 0.10 |
| High Density Lipoprotein, mg/dL (SEM) | 65 (4) | 59 (6) | 0.40 |
| Triglycerides, mg/dL (SEM) | 69 (9) | 85 (19) | 0.40 |
| Hemoglobin A1C, % (SEM) | 5.5 (0.06) | 5.7 (0.3) | 0.40 |
| Insulin, mg/dL (SEM) | 8 (1) | 8 (2) | 0.90 |
| HOMA-IR, US Stanford Units (SEM) | 1.9 (0.2) | 2.1 (0.5) | 0.60 |

### 3.2. Group Differences (Table 2, Figure 3)

Numerical values are summarized in **Table 2**. Results are consistent with the hypothesis of endothelial inflammatory changes in subjects with WMH. Levels of EDE complement effector proteins in the classical and alternative pathways were significantly elevated and complement regulatory proteins were significantly lower in subjects with WMH. After normalizing for EDE concentrations, we found the following factors to have significantly higher levels in subjects with WMH with large effect sizes: C1q (p=.002), C3b (p<.0001), Factor B (p<.0001), Bb (p<.0001), and C5b-9 (p=.0002). EDE levels of the complement regulatory proteins Factor I (p=.02), and CD59 (p<.0001) were significantly lower in subjects with WMH. Results remained significant when adjusting for age in a regression model. The following complement factors and regulatory proteins did not reach statistical significance between groups: Factor D (p=.40), C4b (p=.60), CR1 (p=.60), DAF (p=.10), CD46 (p=.90), and lectin pathway MBL (p=.50).

**Figure 3.**
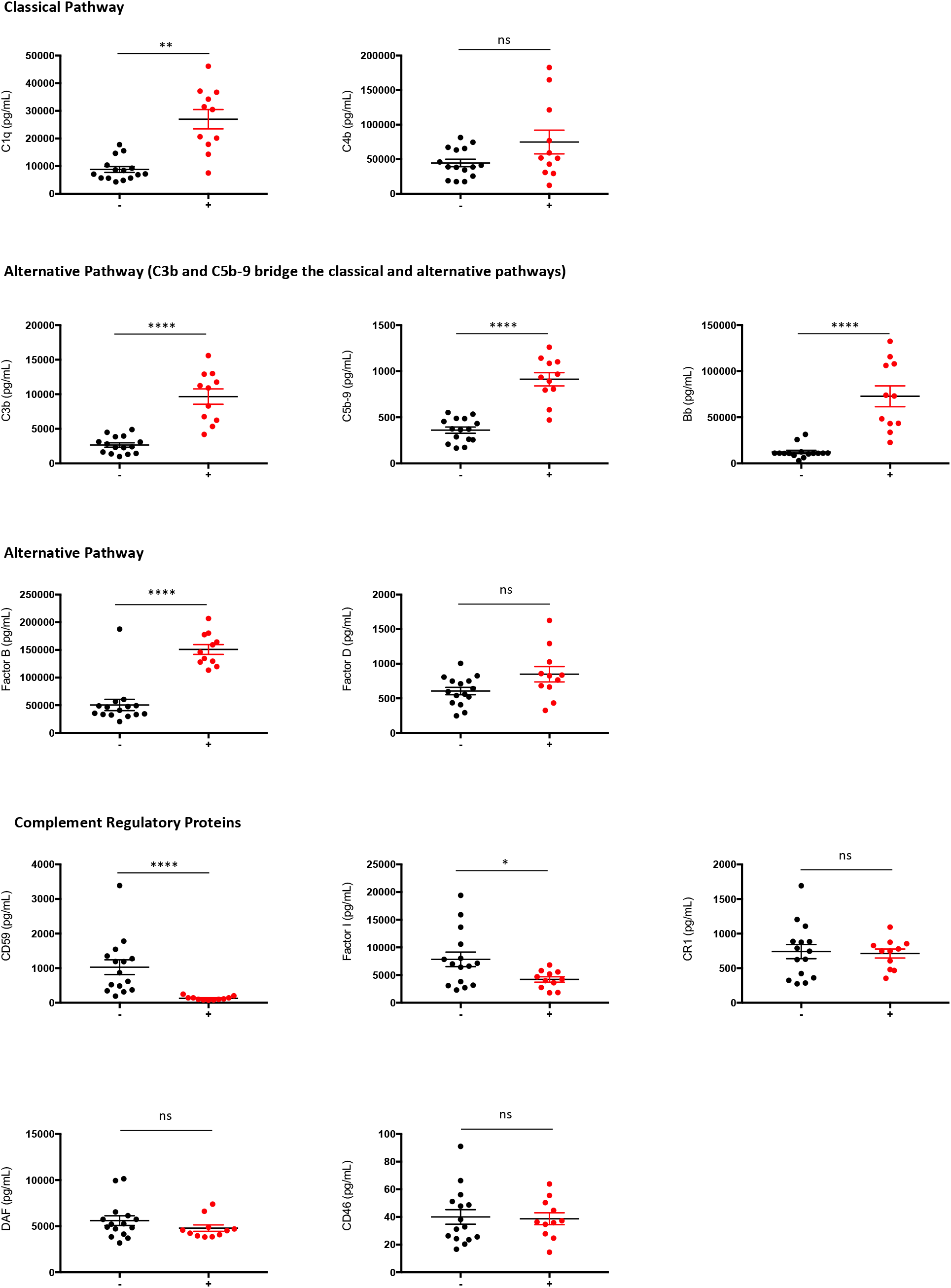
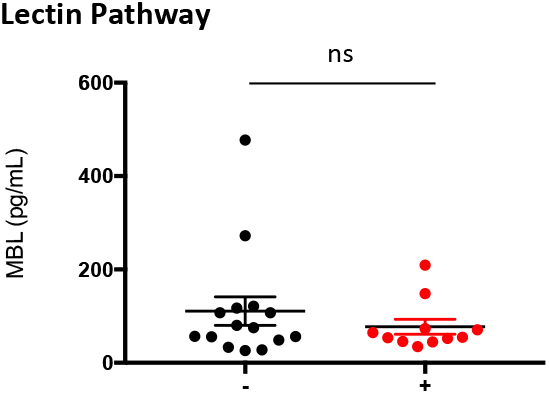
Group comparison of EDE cargo in group of subjects with WMH (+) versus without (−) Complement factors are grouped by pathway. C3b and C5b-9 grouped under alternative pathway are, as depicted in Figure 1, shared across classical, and alternative pathways. Stars depict significance levels from ANCOVA models, controlling for age, detailed in Table 1: ≤.0001 = ****; ≤.001 = ***; ≤.01 = **; ≤.05 = *; > .05 = ns.

**TABLE 2.**
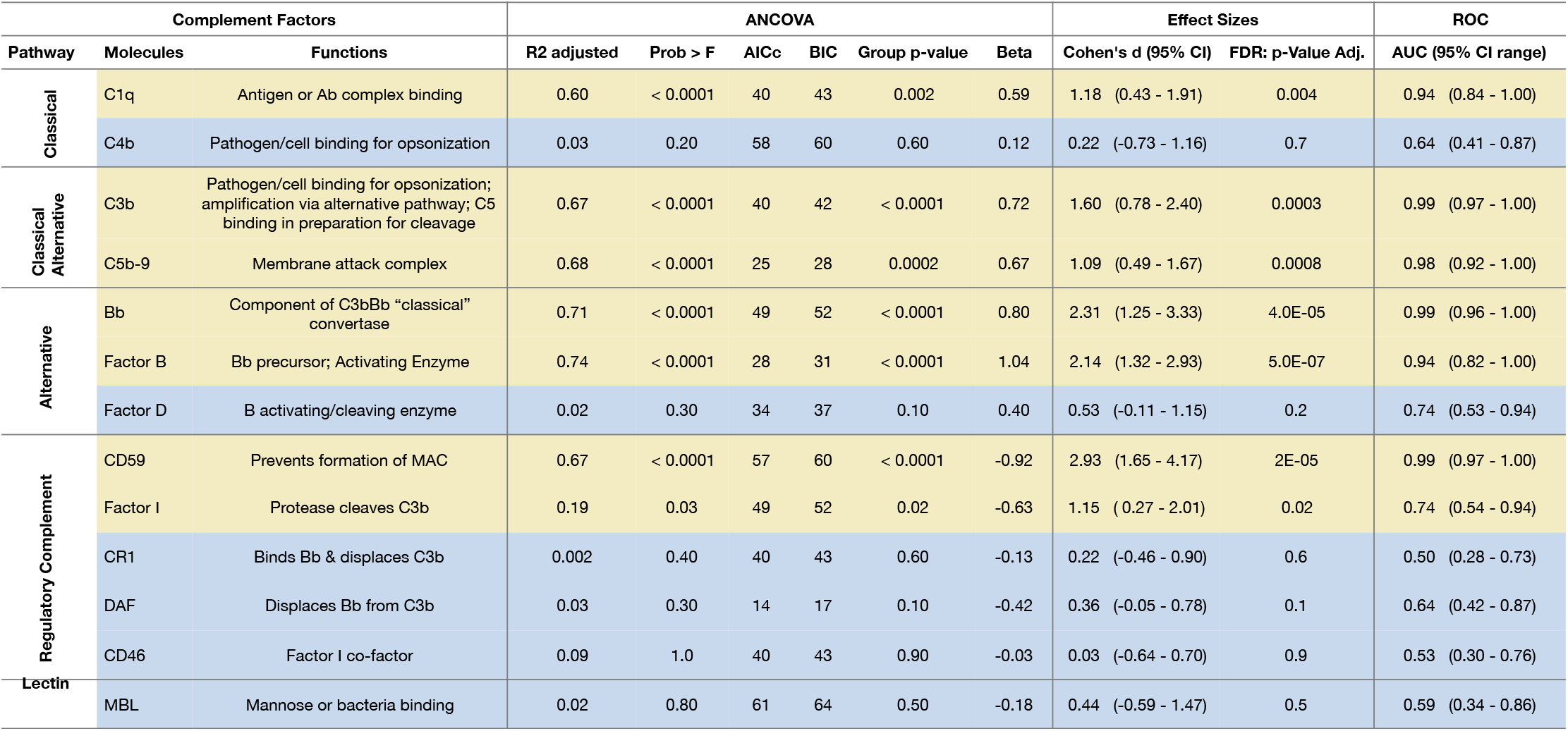
Comparison of EDE complement factor concentrations between groups. This table includes a list of complement factors within pathways and respective functions, and results from statistical models: ANCOVA adjusted for age; Effect sizes (Cohen’s d) based on standard least square models; AUC based from ROC analyses, Acronyms: R^2^ = adjusted for age; CI = confidence interval. Statistically significant values are highlighted in yellow. For ANCOVA, only statistically significant results are provided. P values are rounded to one non-zero digit.

Individual markers had AUC values ranging .74 to .99 (**Table 2**). The greatest group effect sizes were found for C3b, Bb and CD59. For these EDE complement proteins, the likelihood ratio of having significant WMH in comparison to none or mild WMH was found to be 13.64 at sensitivity of 91% (95% CI 59-100) and specificity of 93% (95% CI 68-100). The associated optimal diagnostic accuracy cut off was > 4692 pg/mL for C3b, > 28609 pg/mL for Bb, and < 225 pg/mL for CD59.

### 3.3. Relation with secreted plasma cytokine and chemokine markers (Table 3)

Significant positive associations were found between systemic inflammatory cytokine IL6 and EDE C3b (β=.44; p=.03), C5b-9 (β=.43; p=.03), Bb (β=.40; p=.05), and Factor B (β=.42; p=.04), and negative association with CD59 (β= −.63; p=.0009). We also found a significant association between the endothelial and macrophage expressed adhesion molecule, ICAM1, and EDE C1q (β=.39; p=.05), (C3b (β=.41; p=.04), C5b-9 (β=.44; p=.03), Bb (β=.51; p=.009), and Factor B (β=.47; p=.02), and negative association with CD59 (β= −.59; p=.002).

**TABLE 3.** Results of Linear Models for IL6 and ICAM1 associations with EDE Complement factors. Illustrated in this table are results of standard least square models with: plasma levels of IL6 and ICAM1 as independent variables and EDE cargo as dependent variables.

| Molecules | IL6 |  |  |  |  | ICAM1 |  |  |  |  |
| --- | --- | --- | --- | --- | --- | --- | --- | --- | --- | --- |
|  | R2 adjusted | AICc | BIC | Beta | p-value | R2 adjusted | AICc | BIC | Beta | p-value |
| C1q | 0.10 | 57 | 59 | 0.38 | 0.07 | 0.12 | 57 | 60 | 0.39 | 0.05 |
| C4b | 0.03 | 56 | 58 | 0.18 | 0.40 | 0.006 | 57 | 60 | 0.08 | 0.70 |
| C3b | 0.15 | 60 | 62 | 0.44 | 0.03 | 0.23 | 62 | 65 | 0.41 | 0.04 |
| C5b-9 | 0.15 | 47 | 49 | 0.43 | 0.03 | 0.16 | 47 | 50 | 0.44 | 0.03 |
| Bb | 0.13 | 73 | 75 | 0.40 | 0.05 | 0.23 | 72 | 75 | 0.51 | 0.009 |
| Factor B | 0.14 | 54 | 57 | 0.42 | 0.04 | 0.18 | 55 | 57 | 0.47 | 0.02 |
| Factor D | 0.10 | 30 | 33 | 0.37 | 0.08 | 0.009 | 36 | 39 | 0.09 | 0.70 |
| CD59 | 0.37 | 68 | 70 | -0.63 | 0.0009 | 0.32 | 72 | 74 | -0.59 | 0.002 |
| Factor I | 0.04 | 50 | 53 | -0.28 | 0.20 | 0.70 | 52 | 54 | -0.32 | 0.10 |
| CR1 | 0.0004 | 39 | 41 | 0.03 | 0.90 | 0.03 | 39 | 42 | -0.16 | 0.70 |
| DAF | 0.04 | 14 | 16 | -0.20 | 0.30 | 0.06 | 12 | 15 | -0.32 | 0.10 |
| CD46 | 0.04 | 35 | 38 | -0.20 | 0.38 | 0.05 | 34 | 37 | -0.30 | 0.10 |
| MBL | 0.02 | 55 | 58 | -0.25 | 0.20 | 0.02 | 56 | 58 | -0.24 | 0.30 |

### 3.4. Relation with blood pressure (Table 4)

Significant positive associations were found between an important vascular risk factor, systolic blood pressure, and C1q (β=.46; p=.05) and Bb (β=.58; p=.02) of the classical and alternative pathways, respectively. A significant negative association was found between systolic blood pressure and CD59 (β= −.62; p=.02), a complement regulatory protein.

**TABLE 4.** Relationship between EDE complement proteins and clinical factors. Illustrated in this table are results of standard least square models with: (1) systolic blood pressure as independent variable and complement factors as dependent; (2) complement factors as independent and global volume of white matter hyperintensity as dependent; (3) complement factors as independent and cognitive processing speed (based on time taken to complete the modified Trails test). Age was adjusted for in all models. All p-values are FDR-corrected for multiple comparisons.

| Molecules | Systolic Blood Pressure |  |  |  |  |  | White Matter Hyperintensity |  |  |  |  |  | Processing Speed |  |  |  |  |  |
| --- | --- | --- | --- | --- | --- | --- | --- | --- | --- | --- | --- | --- | --- | --- | --- | --- | --- | --- |
|  | R2 adjusted | Prob > F | AICc | BIC | Beta | p-value | R2 adjusted | Prob > F | AICc | BIC | Beta | p-value | R2 adjusted | Prob > F | AICc | BIC | Beta | p-value |
| C1q | 0.53 | 0.0006 | 38 | 39 | 0.46 | 0.05 | 0.45 | 0.0008 | 155 | 157 | 0.40 | 0.05 | 0.21 | 0.05 | 23 | 25 | 0.59 | 0.05 |
| C4b | 0.06 | 0.60 | 43 | 44 | 0.08 | 0.80 | 0.33 | 0.006 | 159 | 162 | 0.08 | 0.70 | 0.04 | 0.60 | 29 | 31 | 0.08 | 0.70 |
| C3b | 0.32 | 0.01 | 46 | 48 | 0.31 | 0.20 | 0.62 | <.0001 | 146 | 149 | 0.65 | 0.001 | 0.03 | 0.30 | 28 | 29 | 0.33 | 0.20 |
| C5b-9 | 0.33 | 0.01 | 33 | 34 | 0.20 | 0.40 | 0.56 | <.0001 | 149 | 152 | 0.64 | 0.005 | 0.04 | 0.60 | 29 | 31 | 0.13 | 0.70 |
| Bb | 0.52 | 0.0008 | 53 | 54 | 0.58 | 0.02 | 0.47 | 0.0005 | 153 | 156 | 0.40 | 0.03 | 0.37 | 0.006 | 18 | 20 | 0.71 | 0.005 |
| Factor B | 0.08 | 0.20 | 48 | 50 | 0.41 | 0.10 | 0.57 | 0.001 | 149 | 151 | 0.52 | 0.004 | 0.02 | 0.50 | 29 | 30 | 0.17 | 0.50 |
| Factor D | 0.01 | 0.90 | 28 | 30 | 0.02 | 0.90 | 0.37 | 0.003 | 158 | 160 | 0.22 | 0.20 | 0.05 | 0.60 | 29 | 31 | 0.05 | 0.80 |
| CD59 | 0.43 | 0.003 | 58 | 59 | -0.62 | 0.02 | 0.36 | 0.003 | 158 | 161 | -0.21 | 0.20 | 0.14 | 0.10 | 25 | 27 | -0.48 | 0.06 |
| Factor I | 0.10 | 0.16 | 40 | 41 | -0.60 | 0.07 | 0.33 | 0.006 | 159 | 162 | -0.09 | 0.60 | 0.15 | 0.09 | 25 | 26 | -0.44 | 0.10 |
| CR1 | 0.06 | 0.60 | 35 | 36 | -0.17 | 0.60 | 0.32 | 0.007 | 160 | 162 | 0.05 | 0.80 | 0.03 | 0.30 | 28 | 29 | -0.28 | 0.20 |
| DAF | 0.07 | 0.20 | 15 | 17 | -0.49 | 0.09 | 0.37 | 0.003 | 158 | 161 | -0.21 | 0.20 | 0.08 | 0.20 | 26 | 28 | -0.35 | 0.10 |
| CD46 | 0.004 | 1.0 | 36 | 37 | -0.05 | 0.90 | 0.32 | 0.007 | 160 | 162 | 0.01 | 1.0 | 0.001 | 0.40 | 28 | 30 | -0.22 | 0.30 |
| MBL | 0.06 | 0.20 | 50 | 51 | -0.44 | 0.10 | 0.32 | 0.007 | 160 | 162 | 0.006 | 1.0 | 0.02 | 0.30 | 28 | 29 | -0.27 | 0.20 |

### 3.5. Relation with brain imaging (Table 4)

Levels of C3b and C5b-9 involved in both the classical and alternative pathways, and Factor B and Bb of the alternative pathway, were significantly positively associated with global burden of WMH (p-value range .001-.05 detailed in **Table 4**). DAF levels were significantly inversely associated with global volume of WMH (R^2^=.57, β= −42, p=.02). CD46, a key regulatory protein of the end membrane attack complex (**Figure 1**), was significantly associated with total grey matter volumes, controlling for total intracranial volume and age (R^2^=.45, β= .43, p=.03).

### 3.6. Relation with cognitive function (Table 4)

Higher levels of EDE complement proteins C1q (β= .59, p=.05) and Bb (β= .71, p=.005) were associated with significantly lower cognitive function as measured by set-shifting performance speed, controlling for age in all analyses.

## 4. Discussion

We had previously shown high sensitivity and specificity of EDE BBB markers for vascular-mediated brain injury (WMH)^35^. In this study, we extended our previous work into EDE markers of innate inflammation to investigate the hypothesis that endothelial inflammation is involved in white matter injury in functionally normal elders with evidence of WMH on brain imaging. To this end, EDE concentrations of two complement factors in the classical (C1q, and C4b), two in classical and alternative (C3b, and C5b-9), three in the alternative (Factor B, Bb, and Factor D), and one in the lectin pathway (MBL), along with five regulatory proteins (CD59, Factor I, CR1, DAF, and CD46) were compared between groups of subjects with and without evidence of WMH on brain imaging. Overall, our results suggest significant activation of alternative and classical complement pathways in tandem with decreased levels of protective complement regulatory proteins in subjects with WMH.

We demonstrated significant differences for C1q, C3b, C5b-9, Factor B, and Bb, as well as CD59 and Factor I regulatory proteins. All were large effect sizes, particularly for proteins within the alternative pathways Bb, Factor B, and C3b. The largest group differences were found for factors C3b and Bb, such that discriminant function of subjects with WMH versus without reached an AUC of .99 for either factor alone. These findings implicate both the classical and alternative complement pathways, and the resulting opsonizing C3b and the C5b-9 membrane attack complex, in inflammatory endothelial contributions to WMH. The lack of clear difference in EDE MBL levels in subjects with and without WMH suggests that the lectin pathway may not be an important contributor to cerebral endothelial inflammation in functionally normal subjects with WMH. Taken together, the presented findings support involvement of endothelial inflammation in WMH and associations with slowed processing speed in aging. Complement factors involved in innate immune dysfunction are emerging as important molecular contributors to neuronal dysfunction in Alzheimer’s disease^23,36–38^. A prior study has shown activation of alternative and classical complement pathways in astrocytes in AD via astrocytic-derived exosomal analyses^39^. Therefore, the group differences noted in this study likely increase risk of future cognitive decline and Alzheimer’s disease.

In addition to the investigation of group differences in levels of EDE complement markers, we investigated the associations of complement factors with critical vascular risk factors, such as blood pressure, systemic inflammation (plasma IL6 levels), and immune activation (plasma ICAM1 levels), as well as downstream consequences of cerebral vascular disease, such as white matter injury (as a continuous variable), grey matter atrophy, and slowed processing speed. Bb, a key factor in the alternative pathway emerged as significantly associated with both upstream and downstream vascular risk and injury, respectively. Interestingly, a potent inhibitor of the membrane attack complex, CD46, showed a significant positive association with cerebral grey matter volume. CD59 also demonstrated an inverse association with systolic blood pressure. Prior studies have shown changes in EC levels of CD59 in hypoxic states and some microvascular disorders, such as diabetic vasculopathy. It is possible that upregulations of CD59 represent a reactive protective mechanism by endothelial cells. Overall, interactions between inflammatory cytokines and EC complement proteins may enable inflammatory homeostasis or exacerbate a dysfunctional state. Complement binding to ECs results in increased expression of adhesive proteins and production of inflammatory mediators including ICAM1 and IL6, that we measured in this study. Together these changes result in increased vascular permeability and transmigration of PMN leukocytes, resulting in white matter injury and radiographic evidence of WMH.

Since this is the first study to investigate complement activation in endothelial cells of subjects with only radiographic evidence of WMH, we sought to quantify complement factors across all pathways. Overall, abnormal EC-derived exosome cargo levels of complement proteins reflect the nature and extent of dysfunctional complement system homeostasis in ECs and is the basis for the present analytical approach. ECs normally respond to the continuous challenges of inflammatory signals through diverse mechanisms that optimize host defenses and minimize vascular injury. EC exosomes are components of a constitutive system to dispose of cellular “waste” or potentially damaging endogenous inflammatory mediators, in addition to presenting some of them as stimuli to protective blood leukocytes. Complement-mediated injury of ECs in a broad range of vascular diseases is attributable to failures of inflammatory homeostasis that include both enhanced activation of complement effector pathways and diminished complement regulatory elements. Components of activated complement pathways, including C1q, C4b, C3b, or membrane attack complex (MAC) C5b-9 and the anaphylatoxins C5a and C3a are capable of evoking numerous EC inflammatory responses. In addition to representing a means of disposing of unwanted metabolites and proteins, exosomes released from ECs can provide a means of cell-cell communication(ref). Therefore, C3b and C5b-9 packaged in EDEs could potentially get delivered to neurons and display their surface membranes, initiating microglial cytotoxic attacks and neuronal destruction. Moreover, enhanced production of classical and alternative complement factors could be amplified by the relative deficiencies of complement regulatory proteins, also noted in this study (see Figure 1). The main source of complement factors is states of health is the liver. However, it is becoming clear that other cells can also express complement factors. In culture, mammalian ECs have been shown to produce C1, C4, C3, factor B, and factors I among other factors. Of note, although EDE complement factors could be produced by endothelial cells, they could also be sourced elsewhere and simply up taken by ECs. In this study we looked at proteins. RNA-based analyses would clarify sources of transcription of these factors.

In summary, cerebral vascular inflammation is emerging as an important contributor to brain injury and cognitive decline. In addition, higher load of WMH is associated with adverse age-associated cognitive outcomes, such as slowed processing speed and increased risk of AD dementia ^40,41^. In the absence of frank neurodegenerative phenotypes, age-associated WMH is frequently presumed secondary to cerebral vascular endotheliopathies resulting in increased BBB leakage and activation of CNS immune system, leading to reactive gliosis ^40,42–45^. In this model, endothelial cells play a central role. However, a direct *in vivo* interrogation of EC molecular content and testing of specific hypotheses has only just been made possible via EC exosomal isolation techniques. In this proof of concept study, we demonstrate a strong association between endothelial innate inflammation and cerebral white matter injury, a hallmark of small vessel disease, and important contributor to age-associated cognitive slowing, neurodegeneration and dementia. EDE complement factors can provide promising preclinical biomarkers—an avenue that should be pursued further in light of the value of detecting preclinical disease as well as using fluid biomarkers as surrogate outcomes in therapeutic trials targeting endothelial inflammation.

## Acknowledgements

Participant recruitment and data collection for this project was funded by the MarkVCID (UH3NS100608, JH and CD) and Hillblom Aging Network for the Prevention of Age-Associated Cognitive Decline study grants (2140-A-004-NET, JHK). This work was also supported by the NIH-NIA ADRC grants (P50 AG023501, BLM; P30 AG010129, CD).

## Author Contributions

All authors reviewed and edited the manuscript. FE and EG designed the study, performed experiments and interpreted the results, FE performed statistical analyses, figure and table preparation, and wrote the manuscript. DH independently ran all statistical analyses and confirmed the results. MA recruited participants, collected biospecimen and MRI scans. CD and PM processed MRI scans. JK and BM provided funding, resources, and critical input. JH, KC, and AS provided critical input.

## Availability of materials and data

The datasets generated and analyzed during the current study are available from the corresponding author on reasonable request.

